# A rapid and accurate PCR-based method for mating-type determination in the heterothallic oomycete *Plasmopara viticola*, the causal agent of grapevine downy mildew

**DOI:** 10.1101/2025.11.19.689015

**Authors:** C. Couture, I. D. Mazet, Y. Dussert, C. Guyomar, F. Delmotte

## Abstract

*Plasmopara viticola* is a heterothallic diploid oomycete responsible for grapevine downy mildew, one of the most damaging grapevine diseases worldwide. Since no rapid method for mating type characterization of strains is currently available, mating type identification must be performed through crossing assays, which are time-consuming. This study reports the development of two PCR-based molecular markers (MT_indel1 and MT_indel2) based on size variation to distinguish between the mating-type alleles of the oomycete *Plasmopara viticola*. For both markers, P1 strains (Mat-a/Mat-b) exhibited a double-band profile, whereas P2 strains (Mat-a/Mat-a) displayed a single PCR band. The markers were validated on a set of 37 reference strains collected across Europe and showed perfect concordance with mating-type phenotypes determined by crossing tests. They provide a robust, sensitive, and inexpensive DNA-based detection system for mating-type identification in *P. viticola*, offering valuable insights into the pathogen’s reproductive biology and genetic diversity.

## Introduction

*Plasmopara viticola*, the causal agent of grapevine downy mildew, is one of the most devastating pathogens affecting grapevines (*Vitis* spp.) worldwide. Native to North America, this oomycete was introduced into Europe in the late 19th century and has since spread to all major grape-growing regions (Fontaine et al., 2021). *P. viticola* is an obligate biotroph, requiring living host tissue to complete its life cycle, and is characterized by rapid asexual reproduction through sporangia as well as sexual reproduction through oospores.

*P. viticola* is a heterothallic species with two self-incompatible mating types, P1 and P2 (Wong et al., 2001). The pathogen’s ability to evolve and overcome plant resistance, together with its capacity to develop fungicide resistance, poses significant challenges for sustainable vineyard management ((Peressotti et al., 2010), (W. J. Chen et al., 2007), (Paineau et al., 2024), (Delmas et al., 2017), (Wingerter et al., 2021), (Martínez et al., 2025)).

Recent advances in genomics have shed light on the pathogen’s genetic architecture, including the identification of key genomic loci associated with pathogenicity (Paineau et al., 2024), (Dvorak et al., 2025) and mating type (Dussert et al., 2020). The locus controlling the mating-type phenotype is located in a non-recombining, repeat-rich genomic region of approximately 570 kilobases (kb) containing 40 genes. In the P1 mating type, this region is heterozygous (MAT-a/MAT-b), whereas in the P2 type, it is homozygous (MAT-a/MAT-a), indicating dominance of the MAT-b allele.

Traditionally, mating types have been determined through pairing tests between strains, where successful oospore production indicates sexual compatibility. However, this approach is time-consuming, labor-intensive, and requires the availability of grapevine leaves due to the pathogen’s biotrophic lifestyle, as well as access to tester strains of known mating type. The recent availability of *P. viticola* mating-type sequences now enables the development of molecular markers for rapid and reproducible determination of mating type of *P. viticola* strains.

The objective of this study was to develop a set of molecular markers for *P. viticola* mating-type identification. We analyzed whole-genome sequencing data to detect small insertion-deletions (<100 bp), associated with the mating-type phenotype. Indels offer a practical advantage, as they can be readily detected using standard electrophoresis-based methods. Based on this approach, we designed two specific molecular markers capable of discriminating between the MAT-a and MAT-b alleles, thereby distinguishing P1 and P2 strains.

## Material and Methods

### Detection of insertions and deletions in the mating-type locus

We analyzed a set of 54 European *P. viticola* strains (n=26 P1; n=28 P2) that were previously phenotyped and sequenced to identify the mating type locus (Dussert et al., 2020). We investigated genetic variation in the Plvit020 and Plvit030 scaffolds of *P. viticola* genome (Dussert et al., 2020) that include the mating type locus. To this end, we reanalyzed the paired-end sequencing reads of these strains as follows : the preprocessing (trimming of poor-quality reads) step was performed using fastp 2.0 (S. Chen, 2023). The sequencing reads were subsequently aligned to the high-quality reference genome assembly of the INRA-Pv221 strain belonging to the P2 mating type (Dussert et al., 2020) using BWA-MEM 0.1.17 (Li and Durbin, 2009). After filtering and cleaning based on a mapping quality score (MapQ) 20, indel calling was performed using Pindel 0.2.5b9 (Ye et al., 2009) with default parameters and an insert size of 500 bp, for Plvit020 and Plvit030 scaffolds. Pindel indeed detects breakpoints of large deletions, medium sized insertions, inversions, tandem duplications and other structural variants at single-based resolution.

To investigate the relationship between indels presence and mating-type, a correspondance analysis (CA) was performed using FactoMineR 2.8 (Francois Husson; Julie Josse; Sebastien Le;Jeremy Mazet, 2017) on a presence/absence binary table of Pindel results. The presence and absence of structural variants in the selected region across the 54 samples were studied.

The variants with the highest contribution to the first axis and exclusively assigned to one of the mating types were used to select two candidate indels. For each selected indel, we extracted a 400 bp region surrounding the locus of interest on the scaffold from the VCF file generated by Pindel (version v0.2.5b8), using the GATK FastaAlternateReferenceMaker tool (version 4.1.4.1) with the reference genome *Pl. viticola* strain INRA-PV221 (Dussert et al., 2018). Multiple sequence alignment of the region was carried out using Muscle 3.8 (Edgar, 2004) in MEGA (Tamura et al., 2021)(Sup.Figure 1). Primers were designed in the most conserved flanking regions of the candidate indels.

### DNA extraction

Of the 54 strains previously analyzed, a subset of 36 strains along with the reference strain PV2543 of the P1 phenotype were used to validate the markers (n=17 P1; n=20 P2; Sup.Table 1). Those strains were collected from nine countries: the Czech Republic (1), France (3), Germany (11), Hungary (4), Italy (2), Spain (2), and Switzerland (13). The strains were retrieved from cryostorage and propagated on detached grapevine leaves from *Vitis vinifera* cv. Cabernet-Sauvignon plants. The plants were placed in a growth chamber under controlled conditions (22°C, 12-hour light/12-hour dark photoperiod) for 5-7 days. Infected and sporulating leaf discs were subsequently cut and freeze-dried before being ground for DNA extraction.

The DNA of each strain was extracted using a CTAB protocol adapted from(Rogers and Bendich, 1985) with the inclusion of an RNAse A step (Paineau et al., 2024).

### Mating type marker design

Primers pairs spanning sequence variations (InDel) were designed using the Primer3 (Untergasser et al., 2012) web service (https://www.bioinformatics.nl/cgi-bin/primer3plus/primer3plus.cgi). The parameters used were 200 pb size around the indel, 58-62°C primer melting temperature (Tm) and 40-60% primer GC content. For each primer pairs, additional Blastn (version 2.10.0) (Camacho et al., 2009) searches were conducted on a diploid assembly (Paineau et al., 2024) of the *P. viticola* genome to detect potential hybridization in other genomic regions. We also ensured that the designed primers did not match in any region of the grapevine genome in order to avoid potential false positives (diploid assembly from the complete genome of Vitis vinifera *cultivar Chardonnay* cl.04 v2.0) (Minio A., Cochetel N., Figueroa-Balderas R., Cantu D., 2024).

### PCR reaction

PCR amplifications were performed in a 20 μl reaction mixture containing 2 μl of 10x PCR buffer with MgCl2, 0.25 μl 10 mM of a dATP, dGTP, dTTP, and dCTP mix, 0.5 μl 10 μM of the reverse and forward primers (see table 1), 1 μl 10 ng/μl template DNA, and 1 U Taq DNA polymerase. PCR amplification was carried out on an Epi Eppendorf thermocycler with the following program: initial denaturation at 95°C for 3 min, followed by 40 cycles of denaturation at 95°C for 30 s., annealing at 60-61°C (depending on primers) for 30-60 s. (depending on the primers) and elongation at 72°C for 1 min, with a final elongation at 72°C for 10 min. PCR products were stained before loading with VWR® EZ-Vision®, DNA dyes (VWR Chemicals solution) and electrophoresed on a 3% high-resolution agarose gel in 0.5x TBE buffer at a voltage of 5.3 V/cm. Gel imaging was performed using the Gel Imaging Documentation System (Alphaimager 2200, Alpha Innotech, USA) with UV light exposure.

**Table 1.**
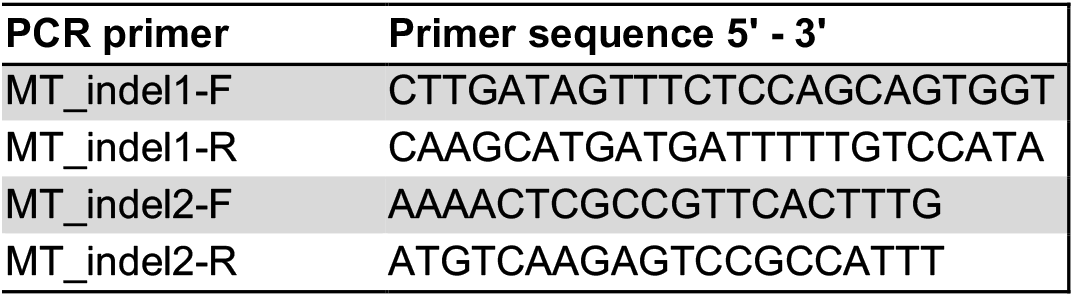
Mating type markers for P. viticola MT_indel1 and MT_indel2.

PCR amplicons were analyzed using the LabChip® GX/GXII Touch™ system (Nucleic Acid Analyzer, PerkinElmer, Waltham, MA, USA) through microfluidic chip electrophoresis, and data were processed with LabChip® GX Reviewer software (version 4.2).

## Results and Discussion

### Identification of indels in the mating-type locus

The objective was to develop genetic markers based on insertion–deletion (indel) polymorphisms capable of distinguishing between the MAT-a and MAT-b alleles, thereby enabling the discrimination of P1 (MAT-a/MAT-b) and P2 (MAT-a/MAT-a) individuals. Pindel detected 76 structural variants in scaffolds Plvit020, Plvit030 and Plvit600, including 68 indels ranging from 1 to 97 bp (Sup.Table 2). Among those indels, 32 were located on Plvit020 and 36 on Plvit030 scaffolds. The first factorial plan of CA of the structural variants table perfectly discriminates strains between the two mating types (Figure 1).

**Figure 1.**
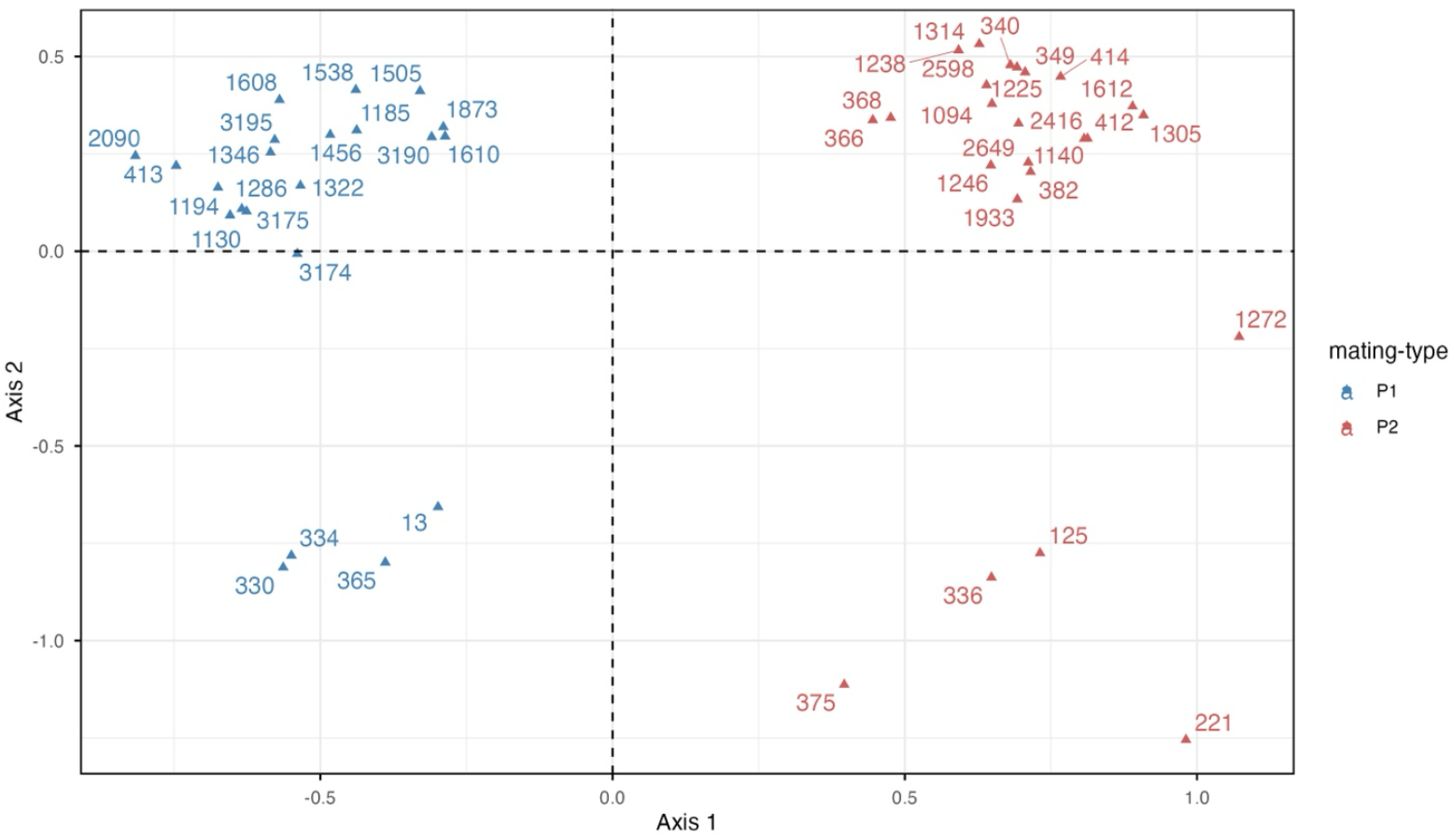
Correspondance Analysis (CA) of P. viticola strains based on the presence/absence of 76 structural variants found in the Plvit020 and Plvit030 scaffolds. Each point represents a strain with positioning reflecting similarity in strutural variant profiles. The axes correspond to the first two CA dimensions, summarizing the main sources of variation in the dataset.

Among the markers significantly associated to axis 1, we retained those consistently present in one mating-type and consistently absent in the other (Sup.Table 3), heterozygous for P1 strains and homozygous for P2 strains, with a minimum coverage allele depth of 10. This resulted in 50 structural variants, including 48 indels. We retained two large indels with a minimum size of 20 bp, in order to enable the development of PCR markers that can be easily visualized on an agarose gel. The first indel was located at 751199, the second one at the position 753867 on the scaffold Plvit030. The definition of primers on these two regions resulted in two candidate markers for the molecular determination of mating type (Table 1).

### Validation of markers

The validation of the two markers was performed using two P1 strains (PV334 and PV13) and two P2 strains (PV412 and PV221). All strains produced results fully consistent with the expected mating-type profiles: P1 strains (MAT-a/MAT-b) displayed two distinct PCR bands, whereas P2 strains (MAT-a/MAT-a) showed a single band.

For the MT_indel1 marker, the two P1 strains produced both the 246 bp and 226 bp fragments, whereas the P2 strains showed only the expected 226 bp fragment (Figure 2). The 226 bp band was more intense in the P2 strains, as this allele is present in two copies.

**Figure 2.**
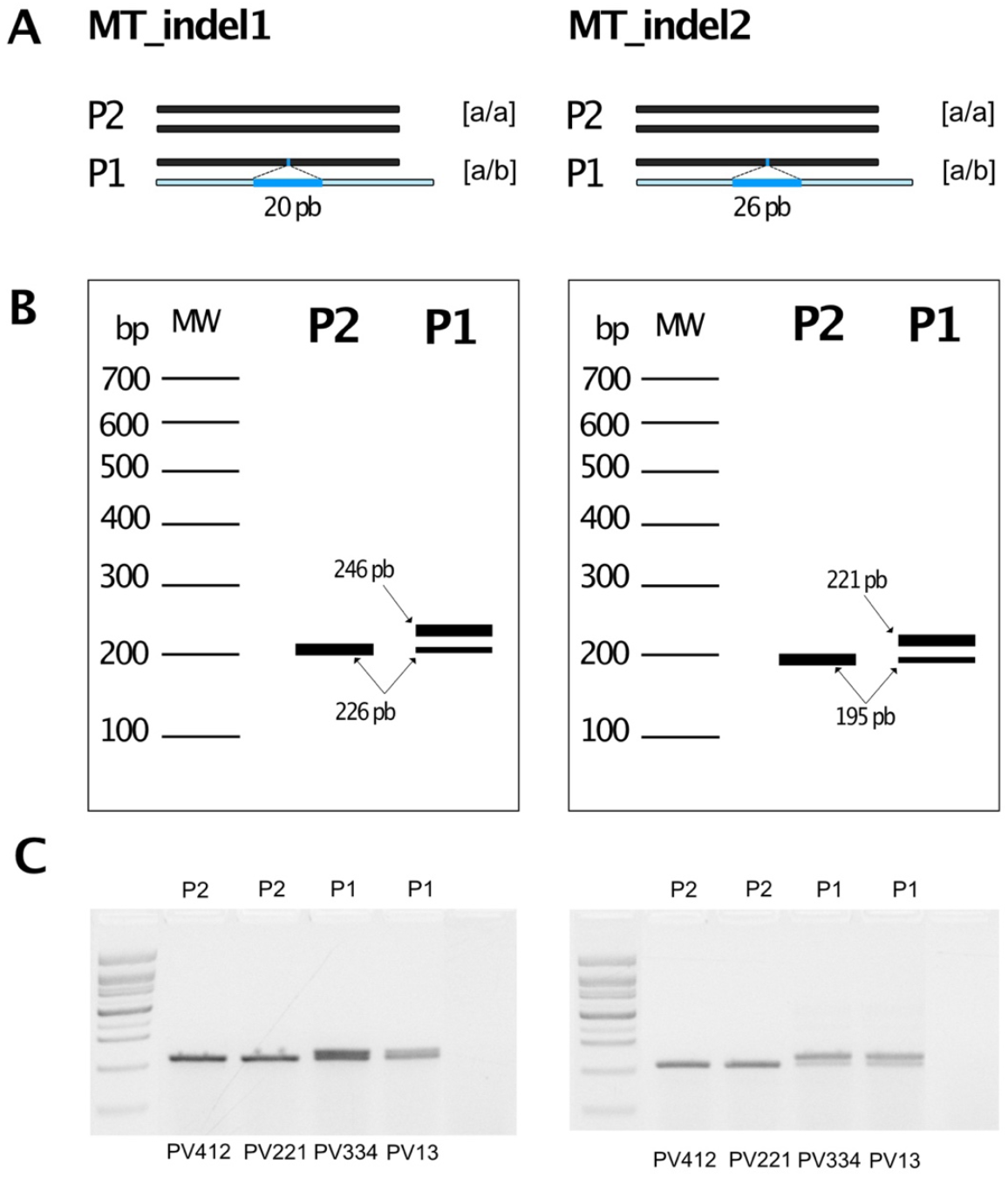
Scoring of P1 and P2 at MT_indel1 and MT_indel2 markers. Both PCR markers are based on the detection of a size difference due to the presence of an insertion in the Mat-b allele relative to the Mat-a allele. Predicted agarose gel electrophoresis patterns for the markers in both P1 and P2 mating types. In P1 individuals, Mat-a and Mat-b alleles are present, producing two bands of distinct sizes. In P2 individuals, only a single allele is present, resulting in a single band. Migration of PCR products on 3% agarose gel electrophoresis amplified with MT_indel1 and MT_indel2 markers.

**Figure 3.**
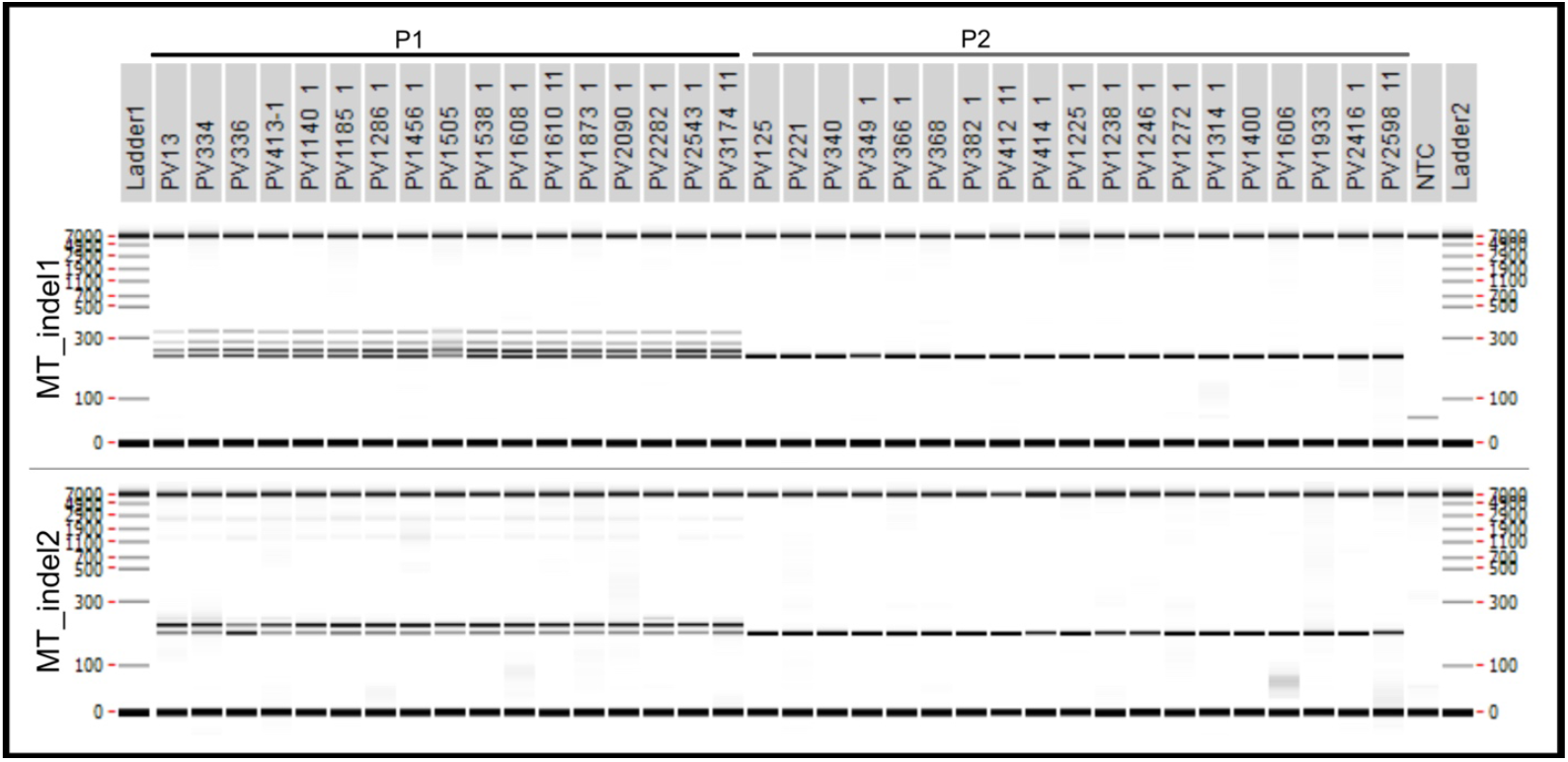
Virtual gels across LabChip® GX/GXII for PCR marker products. MT_indel1 and (226/246 pb) and MT_indel 2 (221/195 pb); (M) – Molecular-weight size marker of 100 bp ladder Extract DNA Ladder 100 pb plus (Ozyme).

For the MT_indel2 marker, the P1 strains yielded both the 221 bp and 195 bp fragments, while the P2 strains displayed only the expected 195 bp fragment (Figure 2). The 221 bp band appeared more intense in the P1 strains for unknown reasons, but this did not interfere with the interpretation of the heterozygous profile.

This analysis also revealed differences in marker performance: the MT_indel1 marker consistently produced stronger signal intensities than MT_indel2, indicating higher PCR efficiency. MT_indel1 generated some low-intensity non-specific bands that were only detectable through microfluidic analysis. These bands may result from repetitive sequences present in other genomic regions, which might not be represented or detectable in the reference genome.

Finally, we tested the markers on a set of 37 reference *P. viticola* strains whose mating type had been previously determined by crossing tests. This confirmed that the markers accurately predict the mating type across a large set of strains representing the population diversity observed in European vineyards. For MT_indel1, P1 strains displayed the expected heterozygous profile, while all P2 strains were homozygous at the locus. For MT_indel2, P1 strains exhibited the heterozygous profile, and P2 strains showed a clear homozygous profile. Therefore, genotyping of the 37 reference strains using the two markers confirmed the association of these molecular markers with the mating-type phenotypes.

The panel of strains used in this study included strains collected from Germany, France, Italy, Hungary, Switzerland, the Czech Republic, Georgia, and Bulgaria, demonstrating the robustness of the markers across a diverse set of strains. Given that *P. viticola* populations in Asia, Australia, and South America originated from Europe (Fontaine et al., 2021), these markers are also expected to be effective for strains from these regions. However, in the pathogen’s center of origin in North America, where genetic diversity is considerably higher, these markers should be further evaluated for their suitability for determinating mating types in local populations.

The development of molecular markers for mating-type identification in *P. viticola* has important implications for future epidemiological studies and disease management. The markers will facilitate the rapid screening of samples to determine their mating type and identify which individuals are capable of producing sexual oospores. They will also enhance our understanding of the reproductive biology and genetic diversity of *P. viticola* by enabling the assessment of mating-type distribution within field populations. This is particularly important, as sexual reproduction influences population genetic diversity and, consequently, the pathogen’s capacity to evolve and adapt to plant resistance or chemical treatments.

In conclusion, the markers provide an inexpensive DNA-based detection system that does not require advanced laboratory equipment. The technique is robust, sensitive, and, most importantly, specific, enabling its use by numerous research laboratories and its easy transfer to extension services working directly with grape growers.

## Acknowledgments

We are gratefull to Fabrice Legeai (INRAE, INRIA) for its assistance in bioinformatics analyses and to Delphine GONZALES (Neurocentre Magendie, Bordeaux University) for tehcnical assistance on electrophoresis.

## Fundings

This study was funded the French National Research Agency (PPR VITAE, grant 20-PCPA-0010) and the Chaire Alexis Millardet (Bordeaux University Fondation).

## Authors’ contributions

CC and IDM conducted the experiments. CG, CC and YD performed the data analysis. CC prepared the initial draft of the manuscript and FD and YD provided feedbacks on the manuscript. FD supervised and led the overall project.

## Notes

### Competing Interest Statement

The authors have declared no competing interest.

